# Deep and accurate detection of m^6^A RNA modifications using miCLIP2 and m6Aboost machine learning

**DOI:** 10.1101/2020.12.20.423675

**Authors:** Nadine Körtel, Cornelia Rücklé, You Zhou, Anke Busch, Peter Hoch-Kraft, FX Reymond Sutandy, Jacob Haase, Mihika Pradhan, Michael Musheev, Dirk Ostareck, Antje Ostareck-Lederer, Christoph Dieterich, Stefan Hüttelmaier, Christof Niehrs, Oliver Rausch, Dan Dominissini, Julian König, Kathi Zarnack

## Abstract

N6-methyladenosine (m^6^A) is the most abundant internal RNA modification in eukaryotic mRNAs and influences many aspects of RNA processing. miCLIP (m^6^A individual-nucleotide resolution UV crosslinking and immunoprecipitation) is an antibody-based approach to map m^6^A sites with single-nucleotide resolution. However, due to broad antibody reactivity, reliable identification of m^6^A sites from miCLIP data remains challenging. Here, we present miCLIP2 in combination with machine learning to significantly improve m^6^A detection. The optimised miCLIP2 results in high-complexity libraries from less input material. Importantly, we established a robust computational pipeline to tackle the inherent issue of false positives in antibody-based m^6^A detection. The analyses are calibrated with *Mettl3* knockout cells to learn the characteristics of m^6^A deposition, including m^6^A sites outside of DRACH motifs. To make our results universally applicable, we trained a machine learning model, m6Aboost, based on the experimental and RNA sequence features. Importantly, m6Aboost allows prediction of genuine m^6^A sites in miCLIP2 data without filtering for DRACH motifs or the need for Mettl3 depletion. Using m6Aboost, we identify thousands of high-confidence m^6^A sites in different murine and human cell lines, which provide a rich resource for future analysis. Collectively, our combined experimental and computational methodology greatly improves m^6^A identification.

**Highlights:**

- miCLIP2 produces complex libraries to map m^6^A RNA modifications
- *Mettl3* KO miCLIP2 allows to identify Mettl3-dependent RNA modification sites
- Machine learning predicts genuine m^6^A sites from human and mouse miCLIP2 data without *Mettl3* KO
- m^6^A modifications occur outside of DRACH motifs and associate with alternative splicing

## INTRODUCTION

The epitranscriptome collectively describes modifications in RNA and has emerged as a crucial and complex mechanism for the post-transcriptional regulation of gene expression. Pervasively occurring in all three kingdoms of life, N6-methyladenosine (m^6^A) is the most prevalent internal modification on mRNA (1, 2). The emerging interest in RNA modifications revealed m^6^A as an essential regulator in almost all aspects of mRNA metabolism and uncovered diverse physiological functions (3–8).

m^6^A is a dynamic modification. It is deposited by writers, recognised by readers and removed by erasers. The writing of m^6^A in mRNA is mainly carried out by a highly conserved, multicomponent methyltransferase complex that catalyses the conversion of adenosine to m^6^A. The methyltransferase like 3 (METTL3) acts as the catalytic active subunit, possessing an S-adenosylmethionine (SAM) binding domain (MTA-70 like domain) with the conserved catalytic DPPW motif (Asp-Pro-Pro-Trp) (9). It installs m^6^A by transferring a methyl group of a SAM donor to targeted adenosines (10). While methyltransferase like 14 (METTL14) is catalytically inactive, it forms a stable heterodimer with METTL3; simultaneously facilitating RNA interaction and increasing the catalytic activity of METTL3 (5,9,11). Additionally, different methyltransferases were identified as m^6^A writers which mainly add m^6^A to U2 and U6 snRNAs, lncRNA or pre-mRNA (12–14). In mRNA, m^6^A enriches in a DRACH ([G/A/U][G>A]m^6^AC[U>A>C]) consensus sequence and occurs in thousands of transcripts, with an average of one to three m^6^A sites per mRNA transcript (15–17). However, only a fraction of DRACH motifs contain an m^6^A modification. Furthermore, m^6^A was found to cluster predominantly within the coding sequence in long internal exons, nearby stop codons and in the 3’ UTR (15, 16).

In order to fully capture and understand the cellular impact of m^6^A, it is essential to precisely locate the modification. Although m^6^A had been identified over four decades ago, only recent fundamental technological breakthroughs allowed transcriptome-wide mapping of m^6^A (15,16,18,19). Antibody-based immunoprecipitation followed by high-throughput sequencing (m^6^A-seq, m^6^A-MeRIP) enabled mapping of m^6^A within a ∼100 nucleotide (nt) window and paved the way to further understand and dissect the cellular and physiological functions of m^6^A (15, 16). Further improvements in 2015 led to an individual-nucleotide resolution UV crosslinking and immunoprecipitation (iCLIP)-based method, called m^6^A iCLIP (miCLIP), which allows the transcriptome-wide mapping of individual m^6^A residues at single-nucleotide resolution (17).

Despite the novel and important insights these epitranscriptomic sequencing methods uncovered, they also suffered several limitations. A critical disadvantage is the required high amount of input material, which makes transcriptome-wide m^6^A detection exclusionary for samples with limited input material. Hence, sequencing low input samples using the aforementioned techniques may lead to over-amplified libraries with a high PCR duplication rate and low complexity. Moreover, it is broadly observed that miCLIP data comprise a lot of background signal due to limited antibody specificity, which makes computational analysis for m^6^A-site identification challenging (20–23).

Here, we present the optimised miCLIP2 protocol, along with the machine-learning-based analysis tool m6Aboost to overcome these limitations. Experimental improvements comprise two separately ligated adapters, two independent cDNA amplification steps and a bead-based size selection (24). These advances result in high-complexity miCLIP2 libraries using less input material at less effort. We performed miCLIP2 in murine embryonic stem cells (mESC), using wild-type (WT) and *Mettl3* knockout (KO) cells to identify peaks that are significantly depleted upon *Mettl3* KO and validated selected m^6^A sites by an orthogonal method. The resulting high-confidence m^6^A sites within DRACH and non-DRACH motifs were used to train a machine learning model, named m6Aboost, which recognises the specific characteristics of m^6^A sites in miCLIP2 data. We applied m6Aboost to multiple miCLIP2 datasets from human and mouse. Thus, our new miCLIP2 protocol in combination with our m6Aboost machine learning model allow to globally predict m^6^A sites in miCLIP2 datasets independently of a *Mettl3* KO.

## MATERIAL AND METHODS

### LC-MS/MS analysis of m^6^A levels

The experiments were performed as described in (25). Ribonucleoside (A, m^6^A) standards, ammonium acetate, and LC/MS grade acetonitrile were purchased from Sigma-Aldrich. ^13^C_9_-A was purchased from Silantes, GmbH (Munich, Germany). ^2^H_3_-m^6^A was obtained from TRC, Inc (Toronto, Canada). All solutions were prepared using ultrapure water (Barnstead GenPure xCAD Plus, Thermo Scientific). 0.1–1 μg of poly(A)+ RNA was degraded to nucleosides with 0.003 U nuclease P1 (Roche), 0.01 U snake venom phosphodiesterase (Worthington), and 0.1 U alkaline phosphatase (Fermentas). Separation of the nucleosides from the digested RNA samples was performed with an Agilent 1290 UHPLC system equipped with RRHD Eclipse Plus C18 (95Å, 2.1 × 50 mm, 1.8 μm, Zorbax, USA) with a gradient of 5 mM ammonium acetate (pH 7, solvent A) and acetonitrile (solvent B). Separations started at a flow rate of 0.4 ml/min and linearly increased to 0.5 ml/min during first 7 min. Then, washing and re-conditioning was done at 0.5 ml/min for an additional 3 min and linearly decrease to 0.4 ml/min during the last minute. The gradients were as follows: solvent B linear increase from 0 to 7% for first 3 min, followed by isocratic elution at 7% solvent B for another 4 min; then switching to 0% solvent B for last 4 min, to recondition the column. Quantitative MS/MS analysis was performed with an Agilent 6490 triple quadrupole mass spectrometer in positive ion mode. Details of the method and instrument settings are described in (26). MRM transitions used in this study were 269.2→137.2 (A), 278.2→171.2 (^13^C_9_-A), 282.1→150.1 (m^6^A), and 285.1→153.1 (^2^H_3_-N6-mrA). Quantification of all samples utilised biological triplicates, and averaged values of m^6^A normalised to A, with the respective standard deviation are shown.

### Cell culture and RNA samples

The HEK293T cell line was cultured in Dulbecco’s Modified Eagle Medium (DMEM, Life Technologies) containing 10% fetal bovine serum (FBS, Life Technologies), 1% L-glutamine (Life Technologies) and 1% penicillin-streptomycin (Life Technologies) at 37°C with 5% CO_2_. All cell lines were monitored for mycoplasma contamination. Anaplastic thyroid carcinoma-derived C643 cells (CLS, RRID:CVCL_5969) were cultured on 15 cm dishes in (DMEM, Thermo Fisher Scientific) supplemented with GlutaMAX (Thermo Fisher Scientific) and 10% FBS at 37°C and 5% CO_2_.

Mouse embryonic stem cells (mESC) with wild-type and *Mettl3* KO genotype were taken from a previous publication (27) and cultured under FBS/LIF conditions as described therein. RAW 264.7 cells (ATCC, Wesel, Germany, TIB-71) were cultured in DMEM (Thermo Fisher Scientific, 12430054) supplemented with 10% heat inactivated FBS (Biochrom, Berlin, Germany, S0613) and 1x penicillin/streptomycin (Thermo Fisher Scientific, 15140-122).

### m^6^A depletion by METTL3 inhibitor treatment

For m^6^A validation in HEK293T cells using SELECT, m^6^A was depleted by using the METTL3 inhibitor STM2457 (STORM Therapeutics)(28). STM2457 was titrated to test for optimal m^6^A depletion quantified by liquid chromatography-tandem mass spectrometry (LC-MS/MS). To this end, HEK293T cells were treated with 2-20 µM STM2457 in DMSO 0.05%-0.2% (v/v) or DMSO alone 0.2% (v/v) as a negative control. After 16 h of treatment, the cells were washed with ice-cold PBS and collected on ice.

### RNA isolation and poly(A) selection

For RNA extraction from HEK293T cells and mESCs, cells were washed in ice-cold PBS and collected on ice for the isolation of total RNA using the RNeasy Plus Mini Kit (Qiagen) following the manufacturer’s recommended protocol. For C643 and RAW 264.7 cells, cells were washed with PBS, and total RNA was extracted using TRIzol reagent (Thermo Fisher Scientific) according to the manufacturer’s instructions. Prior to isolation of poly(A) RNA, total RNA samples were treated with DNase I (New England Biolabs) according to the manufacturer’s protocol, and subsequently cleaned up again by using TRIzol LS (Thermo Fisher Scientific).

For HEK293T and C643 cells, poly(A)+ RNA was extracted using Oligo d(T)_25_ Magnetic Beads using the manufacturer’s recommended protocol (Thermo Fisher Scientific, 61002). Poly(A)+ concentration was measured using Qubit™ RNA HS Assay Kit (Thermo Fisher Scientific). For RAW 264.7 cells, poly(A)+ RNA was extracted by incubating 100 µg total RNA with 200 µl Dynabeads solution (Dynabeads mRNA Direct Purification Kit, Thermo Fisher Scientific, 61012) and purified following manufacturer’s protocols.

The quality of poly(A)+ RNA was ensured using High sensitivity RNA screen tapes for the 2200 Tape station system (Agilent). If a predominant peak for ribosomal RNA was still detectable, an additional round of poly(A) selection was performed, resulting in one round of selection for mESC and RAW 264.7 cells, and two rounds for HEK293T and C643 cells.

### RNA fragmentation

Poly(A)+ RNA was fragmented using RNA fragmentation reagents from Thermo Fisher Scientific. 1 µg of poly(A)+ RNA was filled up to 22 µl with H_2_O for each condition. 1 µl of 0.1-0.4x diluted fragmentation buffer was added (always prepared freshly). The mixture was incubated for 7-12 min at 70°C in thermomixer at 1,100 rpm and put immediately on ice. 1 µl of 0.1 - 0.4x diluted STOP solution was added. The solution was mixed and placed back on ice until use. Time of fragmentation and dilution of fragmentation reaction solutions were optimised prior to miCLIP2 experiments for each new batch of RNA.

### miCLIP2 experiments

All miCLIP2 experiments were performed with rabbit anti-m^6^A antibody purchased from Synaptic Systems (order number 202 003).

#### UV crosslinking and immunoprecipitation

50 µl of protein A Dynabeads (Dynal, 100.02) were magnetically separated, washed two times in 900 µl IP buffer (50 mM Tris, pH 7.4, 100 mM NaCl, 0.05% NP-40) and then resuspended in 50 µl IP buffer and put at 4°C until use. 6 µg of m^6^A antibody was added to the 24 µl of fragmented RNA and rotated for 2 h at 4°C. The IP mixture was placed on a parafilm coated dish and UV irradiated with 2×150 mJ/cm² of UV 254 nm. The mixture was placed back into the tube, another 500 µl of IP buffer and 50 µl of washed protein A beads were added. The mixture was rotated at 4°C for 1 h. The beads were magnetically separated and the supernatant was discarded. The beads were washed two times with high-salt wash (50 mM Tris-HCl, pH 7.4, 1 M NaCl, 1 mM EDTA, 1% Igepal CA-630 (Sigma I8896), 0.1% SDS, 0.5% sodium deoxycholate). The second wash was rotated for at least 1 min at 4°C. Subsequently, the beads were washed two times with PNK buffer (20 mM Tris-HCl, pH 7.4, 10 mM MgCl_2_, 0.2% Tween-20) and resuspended in 1 ml PNK buffer (the samples can be left at 4°C until ready to proceed).

#### 3’ end RNA dephosphorylation

The beads were magnetically separated and resuspended in 20 µl of 3’ end RNA dephosphorylation mixture (4 µl 5x PNK pH 6.5 buffer, 0.5 µl PNK [New England Biolabs; with 3’ phosphatase activity], 0.5 µl RNasin, 15 µl water). The mixture was incubated for 20 min at 37°C in a thermomixer at 1100 rpm. The beads were washed once with PNK buffer, once with high-salt wash (rotate wash for at least 1 min at 4°C) and again washed two times with PNK buffer.

#### L3 DNA linker ligation

The supernatant was magnetically removed and the beads were resuspended in 20 µl of L3 DNA linker ligation mixture (8 µl water, 5 µl 4X ligation buffer, 1 µl RNA ligase [New England Biolabs], 0.5 µl RNasin [N2615, Promega GmbH], 1.5 µl pre-adenylated DNA linker L3-App [20 µM; 5’-/rApp/AGATCGGAAGAGCGGTTCAG/ddC/-3’], 4 µl PEG400 [202398, Sigma]). The mixture was incubated overnight at 16°C at 1,100 rpm in a thermomixer. Subsequently, 500 µl of PNK buffer was added. The beads were washed two times with 1 ml high-salt buffer and two times with 1 ml PNK buffer. After the first wash, the mixture was transferred to a new tube.

#### 5’ end labelling

The beads were magnetically separated and 4 µl of hot PNK mix (0.2 µl PNK [New England Biolabs], 0.4 µl ^32^P-γ-ATP, 0.4 µl 10x PNK buffer [New England Biolabs], 3 µl H_2_O) was added and incubated for 5 min at 37°C in a thermomixer at 1,100 rpm. Next, the supernatant was removed and 20 µl of 1x NuPAGE loading buffer (4x stock was mixed with water and reducing agent and antioxidant was used to avoid potential interference of antibodies) was added to the beads and incubated at 70°C for 5 min.

#### SDS-PAGE and nitrocellulose transfer

The beads were magnetically separated and the eluate was loaded on a 4-12% NuPAGE Bis-Tris gel (Invitrogen). 0.5 l of 1x MOPS running buffer (Invitrogen) was used. Additionally, 5 µl of a pre-stained protein size marker was loaded. The gel was run for 50 min at 180 V. The dye front was cut and discarded as solid radioactive waste. For transferring the protein-RNA complexes to a Protan BA85 Nitrocellulose Membrane, a Novex wet transfer apparatus was used according to the manufacturer’s instructions. The transfer was performed for 1 h at 30 V in 1x transfer buffer with 10% methanol. After the transfer, the membrane was rinsed in 1x PBS buffer. Afterwards it was wrapped in saran wrap and exposed to a Fuji film at 4°C for 30 min, 1 h, or overnight. The film was exposed to a Typhoon phosphoimager.

#### RNA isolation

The protein-RNA complexes were isolated by using the autoradiograph as a mask by cutting the respective regions out of the nitrocellulose membrane. The fragments were placed in a 1.5 ml tube and 10 µl proteinase K (Roche, 03115828001) in 200 µl PK buffer (100 mM Tris-HCl, pH 7.4, 50 mM NaCl, 10 mM EDTA) was added and incubated at 37°C for 20 min at 1100 rpm. 200 µl of PK buffer + 7 M urea (100 mM Tris-HCl pH 7.4, 50 mM NaCl) was added and incubated at 37°C for 20 min at 1,100 rpm. The solution was collected and added together with 400 µl phenol/chloroform (Sigma P3803) to a 2 ml Phase Lock Gel Heavy tube (713-2536, VWR). The mixture was incubated for 5 min at 30°C at 1100 rpm. The phases were separated by spinning for 5 min at 13000 rpm at room temperature. Next, the aqueous layer was transferred into a new tube. Precipitation was performed by addition of 0.75 µl glycoblue (Ambion, 9510), 40 µl 3 M sodium acetate pH 5.5 and addition of 1 ml 100% ethanol. After mixing, the mixture was placed at −20°C overnight. The mixture was spun for 20 min at 15000 rpm at 4°C. After removing the supernatant, the pellet was washed with 0.9 ml 80% ethanol and spun again for 5 min. After removing the supernatant, the pellet was resuspended in 5 µl H_2_O and transferred to a PCR tube.

#### Reverse transcription

RT primers and dNTPs (1 µl primer Rtclip2.0 [5′-GGATCCTGAACCGCT-3’], 0.5 pmol/µl and 1 µl dNTP mix, 10 mM) were added to the resuspended pellet and incubated in a thermocycler (70°C, 5 min, 25°C hold until RT mix is added). After adding the RT mix (7 µl H_2_O, 4 µl 5x RT buffer [Invitrogen], 1 µl 0.1 M DTT, 0.5 µl RNasin, 0.5 µl Superscript III) the mixture was incubated in a thermocycler (25°C, 5 min; 42°C, 20 min; 50°C, 40 min; 80°C, 5 min; 4°C, hold). 1.65 µl of 1 M NaOH was added and incubated at 98°C for 20 min. Subsequently, 20 µl of 1 M HEPES-NaOH pH 7.3 was added. This will eliminate radioactivity from strongly labelled samples after the next step and prevent RNA from interfering with subsequent reactions.

#### Silane clean-up

For bead preparation: 10 µl MyONE Silane beads were magnetically separated per sample and the supernatant was removed. The beads were washed with 500 µl RLT buffer and resuspended in 93 µl RLT buffer. For cDNA binding the beads in 93 µl were added to each sample. After mixing, 111.6 µl of 100% ethanol was added. The mixture was carefully mixed and incubated for 5 min at RT. After incubation, the mixture was again mixed and incubated for 5 min further. After magnetically separating the beads and removing the supernatant, 1 ml of 80% ethanol was added and the mixture was transferred to a new tube. The beads were washed twice in 80% ethanol. The beads were magnetically separated and the supernatant was removed. The tube was briefly mixed in a picoFuge and the remaining supernatant was removed. The beads were air-dried for 5 min at RT. The beads were resuspended in 5 µl H_2_O and incubated for 5 min at RT before performing the on-bead ligation. Radioactivity should be removed. If radioactivity is still detected, continue in hot-lab area.

#### Ligation of 5′ linker to cDNA (on-bead)

The linker was prepared by heating the linker mix (2 µl L##clip2.0 (10 µM stock) 1 μl 100% DMSO) for 2 min at 75°C and keeping it on ice afterwards for > 1 min. The DNA linker L##clip2.0 has the sequence 5′-/5Phos/NNNNXXXXXXNNNNNAGATCGGAAGAGCGTCGTG/3ddC/-3′, where N’s are the 4-nt and 5-nt random nucleotides from the unique molecular identifier (UMI) and X’s are the 6-nt the sample-specific experimental barcode given in **Supplementary Table S1**. After adding the linker mix to the bead containing sample, the ligation mixture (2.0 μl 10x RNA Ligase Buffer [with DTT; New England Biolabs], 0.2 μl 0.1 M ATP, 9.0 μl 50% PEG 8000, 0.3 µl H_2_O, 0.5 μl high conc. RNA Ligase [New England Biolabs]) was pipetted on ice. To ensure homogeneity, the ligation-master-mix was mixed by flicking and spinning it down and was subsequently added with the linker-sample-mix. After vigorous stirring, another 1 µl RNA ligase was added to each sample and mixed by stirring. The mixture was incubated at RT at 1,100 rpm overnight.

#### Silane cleanup of linker-ligated cDNA

Per sample, 5 µl MyONE Silane beads were prepared. The MyONE Silane clean-up was performed as described in the previous Silane clean-up step with following modification: After washing the beads in 500 µl RLT, the beads were resuspended in 60 µl RLT buffer and added to the already bead-containing sample. After the precipitation was performed as previously described, the dried beads were resuspended in 22.5 µl H_2_O.

#### First PCR amplification

The PCR mixture (2.5 µl primer mix 1st PCR [P5Solexa_s, 5′-ACACGACGCTCTTCCGATCT-3′, and P3Solexa_s, 5′-CTGAACCGCTCTTCCGATCT-3], 10 µM each, 25 µl Phusion High Fidelity PCR Master Mix [New England Biolabs, M0531S] was prepared and added to the 22.5 µ of sample from the previous step. A 6-cycle PCR was performed in a thermocycler (98°C, 30 s; 6x [98°C, 10 s; 65°C, 30 s; 72°C, 30 s]; 72°C, 3 min; 16°C, hold).

#### First ProNex size selection

In order to remove primer and primer-dimers, a bead-based size selection was performed prior to preparative PCRs. In addition to the samples, 50 µl of ‘Ultra Low Range Ladder’ (ULR, Thermo Fisher Scientific) will be size selected in parallel to monitor ProNex size selection efficiency. ProNex chemistry was adjusted to RT by keeping it for 30 min at RT. 50 µl of ULR-Phusion mix (1.2 µl ULR Ladder, 28.8 µl H_2_O, 30 µl Phusion PCR mastermix [New England Biolabs] and the samples were mixed with 147.5 µl ProNex chemistry. This is a 1:2.95 v/v ratio of sample:beads. This was optimised in previous experiments (24). The mixture was mixed ten times by pipetting and incubated for 10 min at RT. The sample-bead mixture was placed on a magnetic stand for 2 min and the supernatant was removed. While leaving the bead on the magnetic stand, 200 µl ProNex wash buffer was added to the sample. The buffer was incubated for 60 s before removal. The washes were repeated for a total of 2 washes. After removal of the supernatant, the beads were air-dried for 8-10 min (< 60 min) until cracking starts. The beads were eluted in 23 µl H_2_O. After 5 min of incubation, the mixture was returned to the magnetic stand for 1 min and the supernatant was carefully transferred to a new tube. The size selection efficiency was monitored for the ULR sample on a High Sensitivity D1000 TapeStation Kit. For comparison, the selected and unselected ULR Phusion mix was analysed. The 75-nt/50-nt ladder fragment ratio was compared which should be around 2.5.

#### Optimise PCR amplification

In order to prevent over-amplification of the library, the PCR cycle has to be optimised to a minimum. Therefore, optimise PCR amplification reactions have to be performed for each sample with each 6 and 10 cycles. The PCR mixture (0.5 µl primer mix P5Solexa [5′-AATGATACGGCGACCACCGAGATCTACACTCTTTCCCTACACGACGCTCTTCCGATCT-3′] / P3Solexa [5′-CAAGCAGAAGACGGCATACGAGATCGGTCTCGGCATTCCTGCTGAACCGCTCTTCCGATCT-3′], 10 µM each, 5 µl Phusion High Fidelity PCR Master Mix [New England Biolabs, M0531S], 3.5 µl water) was added to 1 µl of the pre-amplified library. The PCR reaction was performed in a thermocycler (98°C, 30 s; 6 or 10x (98°C, 10 s; 65°C, 30 s; 72°C, 30 s); 72°C, 3 min; 16°C, hold). 2 µl of the amplified library was run on a High Sensitivity D1000 Kit in a TapeStation system. Repeat this step until libraries are seen without over-amplification.

#### Preparative PCR

From previous results of the PCR cycle optimisation, the minimum of PCR cycles was used to amplify ½ of the library. Here, 2.5 times more concentrated cDNA is used, therefore one cycle less is needed than in the preliminary PCR. The PCR mix (8 µl H_2_O, 2 µl primer mix P5Solexa/P3Solexa, 10 µM each, 20 µl Phusion HF Mix [New England Biolabs]) was added to 10 µl cDNA. The PCR was performed in a thermocycler using the same program as in the optimization PCR with the optimised cycle number. 2 µl of the amplified library was run on a High Sensitivity D1000 Kit in a TapeStation system. If the results looked fine, the second half of the library was also amplified and combined with the first half. Finally, the concentration under the peak was determined using TapeStation software, and replicates were combined either in equal molarities or equal volumes.

#### Second size selection by ProNex

Before submitting the samples for sequencing, another round of bead-based size selection was performed to remove residual primers. This ProNex size selection was performed as described above with the following modifications: After ULR preparation, the samples and beads were mixed in a 1:2.4 v/v ratio of sample:beads. This was optimised in previous experiments in (24). After the incubation and washing steps, the dried beads were eluted in 20 µl H_2_O. Again, for comparison the selected and unselected ULR Phusion mix was analysed as described previously. The 100-nt/75-nt ladder fragment ratio should be around 4.5.

### SELECT experiments to validate m^6^A modifications

We used the elongation and ligation-based qPCR amplification method SELECT (29) to independently test for m^6^A modifications at several putative m^6^A sites identified from our miCLIP2 data. Experiments for mESC cells were performed with RNA from mESC WT cells and compared to RNA from mESC *Mettl3* KO cells. Experiments for HEK293T cells were performed with RNA from cells treated with 20 µM METTL3 inhibitor STM2457 (STORM Therapeutics)(28) or DMSO alone as control (see above).

#### Normalisation of input RNA

For *Mettl3* KO or METTL3 inhibitor-treated cell lines, the amount of m^6^A is greatly reduced. Due to m^6^A-mediated RNA degradation or stabilisation processes, absence of m^6^A may influence the abundance of specific transcripts. To ensure usage of same RNA amounts, Qubit (Thermo Fisher Scientific) with Qubit™ RNA HS Assay Kit (Thermo Fisher Scientific) was used to precisely measure RNA concentrations. To ensure usage of equal amounts of transcripts, qPCR experiments were performed for normalisation of input RNA amounts in WT versus m^6^A-depleted cell lines.

#### Elongation and ligation-based qPCR amplification

For the quantitative real-time PCR (qPCR)-based validation of a presumed m^6^A site (termed X site), two primers (Up and Down primer) were designed flanking the site of interest. To precisely measure RNA concentrations before each experiment, Qubit™ RNA HS Assay Kit (Thermo Fisher Scientific) was used. An influence of m^6^A on transcript stability may lead to a difference in transcript abundance upon *Mettl3* KO. Therefore, qPCR for the respective transcript was performed and the amount of total RNA for each SELECT experiment was normalised. To further monitor usage of equal amounts of input material, an Up and Down primer were designed flanking an adjacent nucleotide (termed N site). N sites between X-8 and X+4 were used as input control. According to the previously published SELECT method, 20 ng of poly(A)+ RNA was used per experiment. The RNA was mixed in a total volume of 17 µl in 1xCutSmart buffer containing 40 nM Up primer, 40 nM Down primer and 5 µM dNTPs. The RNA and primers were annealed by incubation in a thermocycler (90°C to 40 °C with a decrease of −10°C after 1 min, then left at 40°C for 6 min). 0.02 U Bst 2.0 DNA polymerase, 0.5 U SplintR ligase and 10 nmol ATP in a volume of 3 µl in 1x CutSmart buffer was added and incubated at 40°C for 20 min. After denaturation at 80°C for 20 min, the mixture was kept at 4°C. Using the Applied Biosystems ViiA7 Real-Time PCR system, qPCR was performed. The 20 µl qPCR reaction mixture contained 2 µl of the final reaction mixture after denaturation, 0.2 nM per qPCR primer, 2x Luminaris HiGreen Lox Rox (Thermo Fisher Scientific) and ddH_2_O. The quantitative qPCR reaction condition was run as follows: 95°C, 5 min; (95°C, 10 s; 60°C, 35 s) x 40 cycles; 95°C, 15 s; 60°C, 1 min; 95°C, 15 s (collect fluorescence at a ramping rate of 0.05°C/s); 4°C hold. qPCR data analysis was performed using QuantStudio Real-Time PCR Software v1.3. All experiments were performed in three technical replicates (separate SELECT reactions). Oligonucleotides used for SELECT are listed in **Supplementary Table S2**.

### RT-PCR quantification of intron retention isoforms

Reverse transcription followed by polymerase chain reaction (RT-PCR) was performed to validate changes in isoform frequencies of selected transcripts (*Ythdc1*, *Mif4gd*) comparing *Mettl3* KO and WT mESCs. Cells were grown on irradiated CF1 mouse embryonic fibroblasts (A34181, Gibco) under normal FCS/LIF conditions, as described before (27). Total RNA was isolated from feeder-depleted mESCs using the RNeasy Plus Kit after removal of genomic DNA with gDNA eliminator columns (Qiagen). Random hexamer primers were used to reverse transcribe 1 µg of total RNA into cDNA using the RevertAid First Strand cDNA Synthesis Kit (Thermo Fisher Scientific) in a thermocycler at 65°C for 5 min, 25°C for 5 min, 42°C for 60 min, 45°C for 10 min, and 70°C for 5 min. Three-primer PCR reactions were performed with One*Taq* DNA Polymerase (New England Biolabs) in a 25 µl reaction, according to the recommended protocol, using 0.5 µl cDNA as template, a shared forward primer located in the upstream exon and two isoform-specific reverse primers in the intron (IR) and the downstream exon (spliced), respectively. All three primers were used in a final concentration of 200 nM each, rendering the shared primer as a rate-limiting factor in the reaction. Primer sequences were: Ythdc1_shared (5’-CCATCCCGTCGAGAACCAG-3’), Ythdc1_IR (5’-CCAACGTGACCATGTGAAATCC-3’), Ythdc1_exonic (5’-TGGTCTCTGGTGAAACTCAGG-3’), Mif4gd_shared (5’-CCTGAGAGTCTGAGCAGGGA-3’), Mif4gd_IR (5’-AAGCCTTGGCCTCTATGTGC-3’) and Mif4gd_exonic (5’-AGCCGTCCCGGATTAGGATA-3’). The PCR reaction was carried out in a thermocycler at 94 °C for 30 s, 30 cycles of [94°C for 30 s, 55°C (*Mif4gd*) or 54°C (*Ythdc1*) for 1 min, 68°C for 1 min] and final extension at 68°C for 5 min. PCR products were analysed by capillary gel electrophoresis on the TapeStation 2200 system using D1000 Screen Tapes (Agilent) according to the manufacturer’s recommendations. Band intensities were quantified using the TapeStation Analysis Software and frequency was calculated as the relative proportion of IR and spliced transcript abundance.

### miCLIP2 read processing

Multiplexed miCLIP2 libraries were sequenced as 91-nt or 92-nt single-end reads on an Illumina NextSeq500 sequencing system including a 6-nt sample barcode as well as 5-nt+4-nt unique molecular identifiers (UMIs).

Initial data processing was done as described in Chapters 3 and 4.1 of (30) for iCLIP data. In short, after checking the sequencing qualities with FastQC (v0.11.8) (https://www.bioinformatics.babraham.ac.uk/projects/fastqc/) and filtering reads based on sequencing qualities (Phred score) of the barcode region (FASTX-Toolkit v0.0.14) (http://hannonlab.cshl.edu/fastx_toolkit/), seqtk v1.3 (https://github.com/lh3/seqtk/), reads were de-multiplexed based on the experimental barcode (positions 6 to 11 of the reads) and adapter sequences were removed from the read ends (Flexbar v3.4.0) (31). UMIs were trimmed as well and added to the read names. Reads shorter than 15 nt were removed from further analysis. Individual samples were then mapped to the respective genome (assembly version GRCh38.p12 for all human samples, GRCm38.p6 for all mouse samples) and its annotation (GENCODE release 31 for all human samples, GENCODE release M23 for all mouse samples) (32) using STAR (v2.7.3a) (33). When running STAR (with parameter --outSAMattributes All), up to 4% mismatches were allowed per read, soft-clipping was prohibited on the 5’ end of reads and only uniquely mapping reads were kept for further analysis. Following mapping, sorted BAM files were indexed (SAMtools v1.9) (34) and duplicate reads were removed (UMI-tools v1.0.0) (35). Reads were defined duplicates if their 5’ ends map to the same position and strand in the genome and they have identical UMIs.

After removing duplicates, all mutations found in reads were extracted using the Perl script parseAlignment.pl of the CLIP Tool Kit (CTK, v1.1.3) (36). The list of all found mutations specifies the mutations, their locations in the genome as well as the names of the reads in which they were found. The list was filtered for C-to-T mutations using basic Bash commands and kept in BED file format as described in (37). Based on the filtered list of C-to-T mutations, de-duplicated reads were separated into two BAM files holding reads with and without C-to-T mutation, respectively, using SAMtools and basic Bash commands. The BAM file of reads without C-to-T mutation was transformed to a BED file using bedtools bamtobed (BEDTools v2.27.1) (38) and considering only the 5’ mapping position of each read. Afterwards, the BED file was sorted and summarised to strand-specific bedGraph files which were shifted by one base pair upstream (since this nucleotide is considered as the cross-linked nucleotide) using bedtools genomecov (BEDTools v2.27.1). Similarly, the BED files of C-to-T mutations were also sorted and summarised to strand-specific bedGraph files using bedtools genomecov. Finally, all bedGraph files were transformed to bigWig track files using bedGraphToBigWig of the UCSC tool suite (v365) (39). Reads containing C-to-T transitions were filtered out using parseAlignment.pl from the CTK package (36) as described in (37).

The code for miCLIP2 data processing as described here is available from two recent data analysis publications (30, 37).

### Peak calling, transcript assignment and relative signal strength

Bam files with reads without C-to-T mutation were used for peak calling with PureCLIP (v1.3.1) (40) individually on each replicate for each condition. PureCLIP significant sites per replicate were then filtered for presence in at least two replicates for a given condition (PureCLIP peaks in **Supplementary Table S1**). For assigning a host gene to each PureCLIP peak, transcript annotations were taken from GENCODE (release 31, GRCh38.p12 for human and release M23, mm10 for mouse), and filtered for a transcript support level ≤ 3 and support level ≤ 2. For overlapping transcripts, the longest annotation was chosen. We next assigned the miCLIP2 peaks to the transcripts.

In order to calculate the relative signal strengths of all peaks within a transcript, we calculated the mean number of truncation events for all peaks in the same transcript. Then, we divided the individual truncation read number of each peak by the mean of the peak strength in the corresponding transcript, leading to a value representing the relative peak strength.

### Differential methylation analysis to identify Mettl3-dependent m^6^A sites

Similar to iCLIP, the miCLIP2 signal is strongly influenced by the underlying transcript abundance (41, 42). Therefore, when applying DESeq2 (43) collectively to all peaks (*one-run*), any change of transcript abundance will lead to incorrect fold change and FDR estimations, resulting in false positive calls in down-regulated genes. We tested four different approaches to overcome this, namely separately running DESeq2 on peaks of individual genes (*gene-wise*) or groups of genes with similar abundance change (*bin-based*), by building a combined DESeq2 model on peak signals and transcript counts using interaction terms (*2-factor*) as well as by using DEXSeq (*dexseq-run*) (44) instead of DESeq2. The different approaches are explained in more detail in the **Supplementary Material**. The best performance was seen for the *bin-based* approach, which was used for all following analyses.

### Training and evaluation of the machine learning model m6Aboost

Based on the log2-transformed fold change (log2FC) and the false discovery rate (FDR) from the *bin-based* differential methylation analysis between WT and *Mettl3* KO cells, we used peaks at A to compile a positive (log2FC < 0, FDR ≤ 0.01; n=11,707) and negative (log2FC ≥ 0, FDR > 0.5; n=42,090) set. Both were combined and then randomly split into a training set (80%) and an independent test set (20%). We then extracted 27 features, including the nucleotide sequence in a 21-nt window around the central A, the transcript region as well as the relative signal strength (log_2_) and the number of associated C-to-T transitions (log_2_). We initially tested three different machine learning algorithms (AdaBoost, support vector machine [SVM], random forest) and evaluated their performance based on precision-recall curves and area under the curve (AUC) as well as by comparing F1-score, Matthews correlation coefficient (MCC), precision, accuracy, sensitivity and specificity on the independent test set. Based on these measures, we selected the AdaBoost-based predictor, which we named m6Aboost (see **Supplementary Material**, Section B for details).

### RNA-seq read processing

RNA sequencing (RNA-seq) libraries were sequenced on an Illumina NextSeq500 as 84-nt single-end reads, yielding 31-35 million reads per sample. Basic sequencing quality checks were applied to all reads using FastQC (v0.11.8) (https://www.bioinformatics.babraham.ac.uk/projects/fastqc/). Reads were mapped to the mouse genome (assembly version GRCm38.p6) and its annotation based on GENCODE release M23 using STAR (v2.6.1b) (33). When running STAR, up to 4% mismatches were allowed per read and only one location was kept for multi-mapping reads. Coverage tracks for visualisation were obtained by merging bam files for each condition using samtools (v1.11). Coverage was calculated with bamCoverage (v3.5.0) from the deepTools suite (45) using RPGC normalisation and –effectiveGenomeSize calculated by ucsc-facount (v377).

For differential gene expression analysis, mapped reads were counted with htseq-count (v0.12.4, -s reverse) (46) into gene annotation based on GENCODE release M23. Differential expression analysis was performed with DESeq2 (v1.30.0) (43) using the method “apeglm” for shrinkage of log_2_-transformed fold changes.

Intron retention (IR) analysis was done with IRFinder (v1.3.0) (47) using built-in script analysisWithLowReplicates.pl for differential analysis (48). We adapted some built-in filtering steps by overwriting line 179 of analysisWithLowReplicates.pl into:

*my $ok = ($pA[8] > 0 || $pB[8] > 0) && ($pA[19] > 0 || $pB[19] > 0) && separatedAB(\@repsIR, $repsA, $repsB);*

and line 186 into:

*if (($pA[8] > 0 || max($pA[16],$pA[17]) > 0) && ($pB[8] > 0 || max($pB[16],$pB[17]) > 0)) {*

For downstream analysis, IR events were filtered for IRratio ≥ 0.03 in at least one condition and mean IntronDepth ≥ 3. *P* values were corrected using Benjamini-Hochberg adjustment.

### Overlap with MAZTER-seq

Processed MAZTER-seq data from (21) were downloaded from Gene Expression Omnibus (GEO) via accession number GSE122956. The m^6^A sites therein were filtered for a difference in MazF cleavage efficiency > 0.1 between WT and *Mettl3* KO, yielding a total of 580 reliably identified m^6^A sites from mESC cells. 200 of these (34.5%) overlapped at single-nucleotide resolution with the 4,464 predicted m^6^A sites at ACA from our mESC miCLIP2 data.

### YTHDF1 iCLIP processing and overlap with predicted m^6^A sites

YTHDF1 iCLIP reads were quality filtered and processed as in Busch et al, 2020 (30), used tools versions are as described above for miCLIP2. For peak calling with PureCLIP (40) reads from the four replicates were merged. Resulting peaks were filtered to be present in at least two out of four replicates. To generate binding sites, peaks closer than 4 nt were merged, allowing no overlapping binding sites. Finally, binding sites were centred at the position with the highest truncation read number as described in (30). All predicted m^6^A sites were aligned and spanned with a 21-nt window to count the presence of YTHDF1 binding sites in that area.

## RESULTS

### The miCLIP2 protocol allows profiling of m^6^A RNA modifications

In order to allow for deep m^6^A profiling, we combined the miCLIP procedure with our recently optimised iCLIP2 protocol, termed miCLIP2 (**Figure 1A**) (17, 24). Experiments were performed with poly(A)+ RNA from mouse embryonic stem cells (mESCs). We first performed two consecutive rounds of poly(A)+ RNA enrichment for total RNA samples (**Supplementary Figure S1A**) and optimised the RNA fragmentation time required for each sample (**Supplementary Figure S1B**). The RNA was then incubated with an m^6^A-specific antibody (Synaptic Systems), which was previously shown to yield highest truncation efficiency in miCLIP experiments (**Figure 1A**) (17). After optimising UV irradiation (254 nm twice with 150 mJ/cm² strength; **Supplementary Figure S1C**), crosslinked antibody-RNA complexes were immunoprecipitated using protein A beads. Co-purified RNAs were 3’dephosphorylated with T4 polynucleotide kinase (PNK) prior to first adapter ligation (L3-APP) and radioactive labelling. After SDS-PAGE gel and transfer, the respective nitrocellulose membrane fragment was excised (**Supplementary Figure S1D**). Transferred RNA was recovered by proteinase K treatment, leaving a polypeptide at the crosslinking site. Reverse transcription generally truncates at this polypeptide thus encoding the positional information about m^6^A sites within resulting cDNA fragments (17, 49). The residual readthrough events usually incorporate C-to-T transitions (17), which provide additional confidence for truncation-identified crosslink sites (see below). After bead-based clean-up and second linker ligation, a pre-amplification PCR (6 cycles) was employed to minimise loss of information by potential material loss in the following steps. This was followed by size selection to remove primer dimers and a second PCR which was optimized for a minimal number of PCR cycles to obtain sufficient material for sequencing (here 11 cycles). After a second size selection to remove remaining primers, the library was subjected to high-throughput sequencing (**Supplementary Figure S1E**).

**Figure 1.**
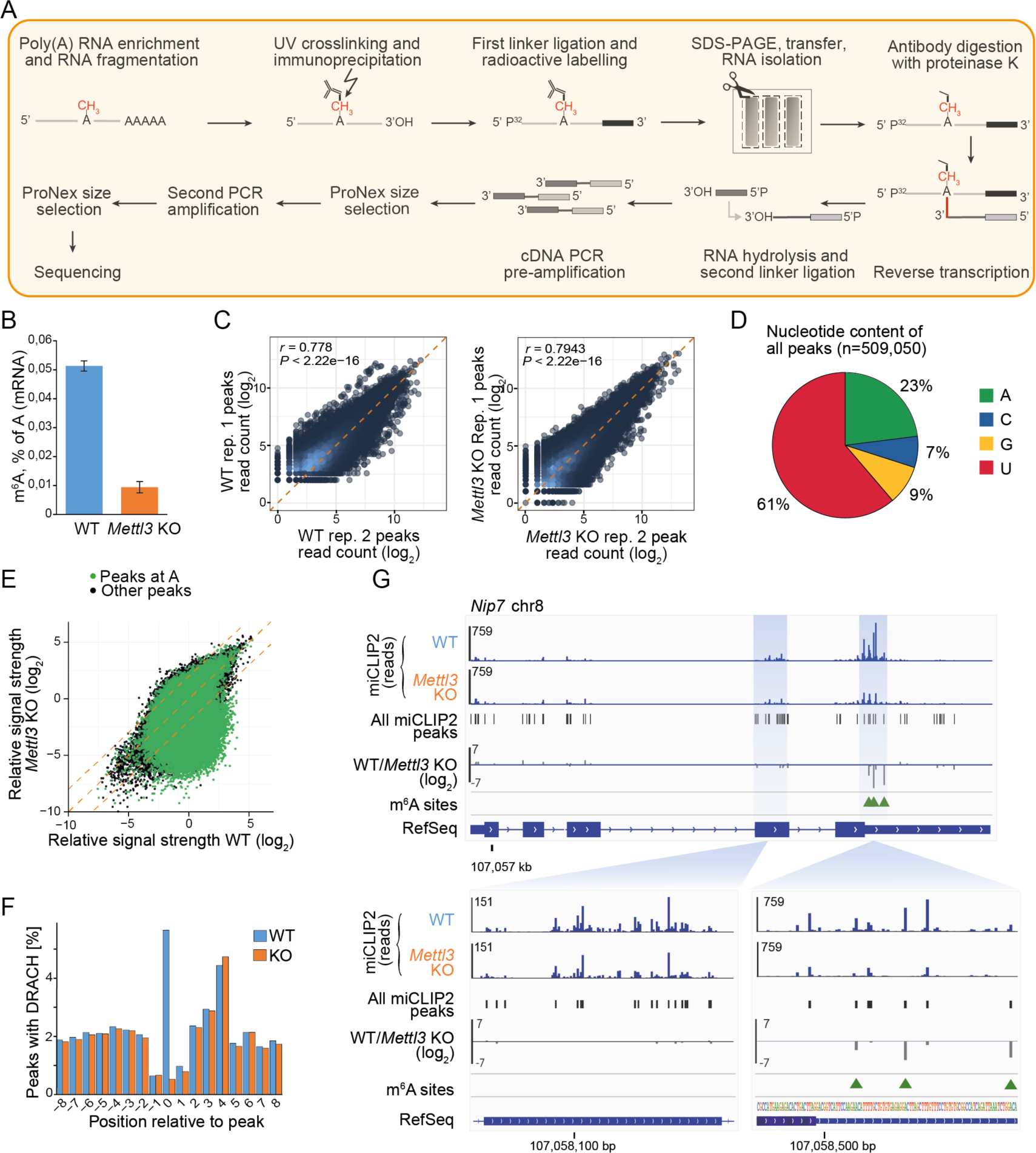
The optimised miCLIP2 protocol produces high complexity libraries with high reproducibility. ***A.*** An overview of the miCLIP2 protocol. ***B.*** mESC *Mettl3* KO cells show a significant depletion of m^6^A on mRNAs. m^6^A levels measured by liquid chromatography-tandem mass spectrometry (LC-MS/MS) for poly(A)+ RNA from WT and *Mettl3* KO mESCs. Quantification of m^6^A as percent of A in mRNA. Error bars indicate standard deviation of mean (s.d.m.), n = 3. ***C.*** miCLIP2 data are highly reproducible between replicates. Pairwise comparison of the miCLIP2 truncation reads within peaks from two miCLIP2 replicates from WT and *Mettl3* KO mESCs. Pearson correlation coefficients (*r*) and associated *P* values are given. Additional replicates are shown in **Supplementary Figure S2A**. ***D.*** Most peaks are located at uridines and adenines. Pie chart representing the nucleotide distribution of all miCLIP2 WT peaks. ***E.*** The majority of peaks are unchanged in a *Mettl3* KO miCLIP2 experiment, indicating high background signal. Scatterplot of the log_2_-transformed relative signal strength (corrected for transcript abundance) of all miCLIP2 peaks in WT and *Mettl3* KO mESC. Peaks located at an A are highlighted in green. Dotted lines indicate diagonal and 4-fold change. ***F.*** DRACH motifs are enriched at miCLIP2 WT peaks. Metaprofile of DRACH motifs around aligned miCLIP2 peaks (position 0). Percentage of DRACH motifs (counted at position of A within DRACH) around the miCLIP2 peaks of WT and *Mettl3* KO mESCs are shown. ***G.*** *Mettl3* KO miCLIP2 signal is reduced at specific positions in the *Nip7* 3’ UTR. Genome browser view of miCLIP2 data (blue) from WT and *Mettl3* KO mESCs and fold change between conditions (grey). Identified miCLIP2 peaks (black bars) and m6Aboost-predicted m^6^A sites (green arrowheads) are given. Zoom-ins (bottom) show more detailed views of an exonic region without m^6^A sites and a 3’ UTR region with three m^6^A sites.

### The majority of miCLIP2 peaks are not sensitive to *Mettl3* KO

In order to test whether miCLIP2 peaks are dependent on Mettl3, we performed miCLIP2 experiments (n=3 replicates) from wild-type (WT) as well as *Mettl3* knockout (KO) mESCs. The latter lack the primary m^6^A methyltransferases Mettl3 and hence, lost most of m^6^A mRNA methylation (**Figure 1B**) (27, 50). Reads with C-to-T transitions (6%) were removed for later usage (**Supplementary Table S1**). The remaining reads corresponded to a total of 261 million putative truncation events (**Supplementary Table S1**). Peak calling on the data from WT mESC cells identified more than 500,000 peaks that exceeded the local background signal (peaks on all samples are reported in **Supplementary Table S1**). The number of truncation events in called peaks were highly reproducible between replicates (**Figure 1C and Supplementary Figure S2A**). To allow for quantitative comparisons between transcripts, we calculated the relative signal strength of all peaks, which was independent of the underlying transcript abundance (see Methods; **Supplementary Figure S2B**).

Analysis of the underlying sequence showed that most peaks resided on thymidine rather than adenosine and only 25% of these adenosines were part of a DRACH motif (**Figure 1D-G**), reflecting UV crosslinking biases and limited antibody specificity as reported previously (20, 21). Nevertheless, the strongest peaks frequently coincided with AC and were located precisely on the A nucleotide (**Supplementary Figure S2C**). We noted an additional enrichment of AC downstream of the peaks. However, these particular peaks did not harbour a DRACH motif and their signal was not reduced in the *Mettl3* KO, indicating that they are part of the unspecific background signal of the employed antibody or m^6^A sites independent of Mettl3 (**Supplementary Figure S2C**). Importantly, peaks at A, AC and DRACH motifs were specifically lost in the *Mettl3* KO, supporting that miCLIP2 detects Mettl3-dependent m^6^A modifications (**Figure 1E-G and Supplementary Figure S2D**). In addition to the putative m^6^A sites, we observed an accumulation of miCLIP2 truncation events at transcript start sites which did not respond to the *Mettl3* KO (**Supplementary Figure S2E and F**). This likely reflects the related RNA modification N6,2’-O-dimethyladenosine (m^6^Am) which is known to reside at 5’ cap structures and is also recognised by the m^6^A-specific antibody (17). Overall, the high amount of nonspecific background and cross-reactivity in the miCLIP2 data required more precise measures to define true Mettl3-dependent m^6^A sites.

### Differential methylation analysis detects Mettl3-dependent m^6^A sites at DRACH and non-DRACH motifs

In order to learn about the features of genuine m^6^A sites in the miCLIP2 data, we sought to extract all miCLIP2 peaks that significantly changed in the *Mettl3* KO mESCs. However, changes at individual peaks were overshadowed by massive shifts in gene expression in *Mettl3* KO cells, with more than 2,809 genes altered at least 2-fold in comparison to WT mESCs (false discovery rate [FDR] ≤ 0.01; **Figure 2A**). These massive shifts in the underlying transcript abundances meant that miCLIP2 read counts at individual peaks could not be compared directly. In order to overcome this shortcoming, we tested several strategies for differential methylation analysis to account for the substantial gene expression changes in the *Mettl3* KO cells (see **Supplementary Material**, Section A). Best performance was achieved with the *bin-based* approach, in which genes were stratified according to their expression change upon *Mettl3* KO (**Figure 2B and Supplementary Figure S3A-C**). All miCLIP2 peaks within the genes of same bin, i.e., with a similar change in gene expression, were then tested collectively using DESeq2 (43) (see **Supplementary Material**, Section A). As expected, the changing peaks almost exclusively showed a loss of miCLIP2 signal in the *Mettl3* KO (**Figure 2C**), and 85.3% of these downregulated peaks were located at A (**Figure 2D**), supporting that our differential methylation analysis enriched for m^6^A sites. From these, we compiled a stringent set of 11,707 sites at A with reduced signal in the *Mettl3* KO (log_2_-transformed fold change [log2FC] < 0, FDR ≤ 0.01), which served as ‘positive set’ of true m^6^A sites in the following analyses (see **Supplementary Material**, Section A). As previously described, the positive sites accumulated nearby stop codons and in 3’ UTRs, and the underlying sequences resembled the DRACH motif (16, 51) (**Figure 2E and F**), supporting that they indeed represented Mettl3-dependent m^6^A sites. For comparison, we selected a ‘negative set’ of 42,090 peaks that were also located at A but unchanged or even mildly increased upon *Mettl3* KO (log2FC ≥ 0, FDR > 0.5) and hence represented the nonspecific background in the data.

**Figure 2.**
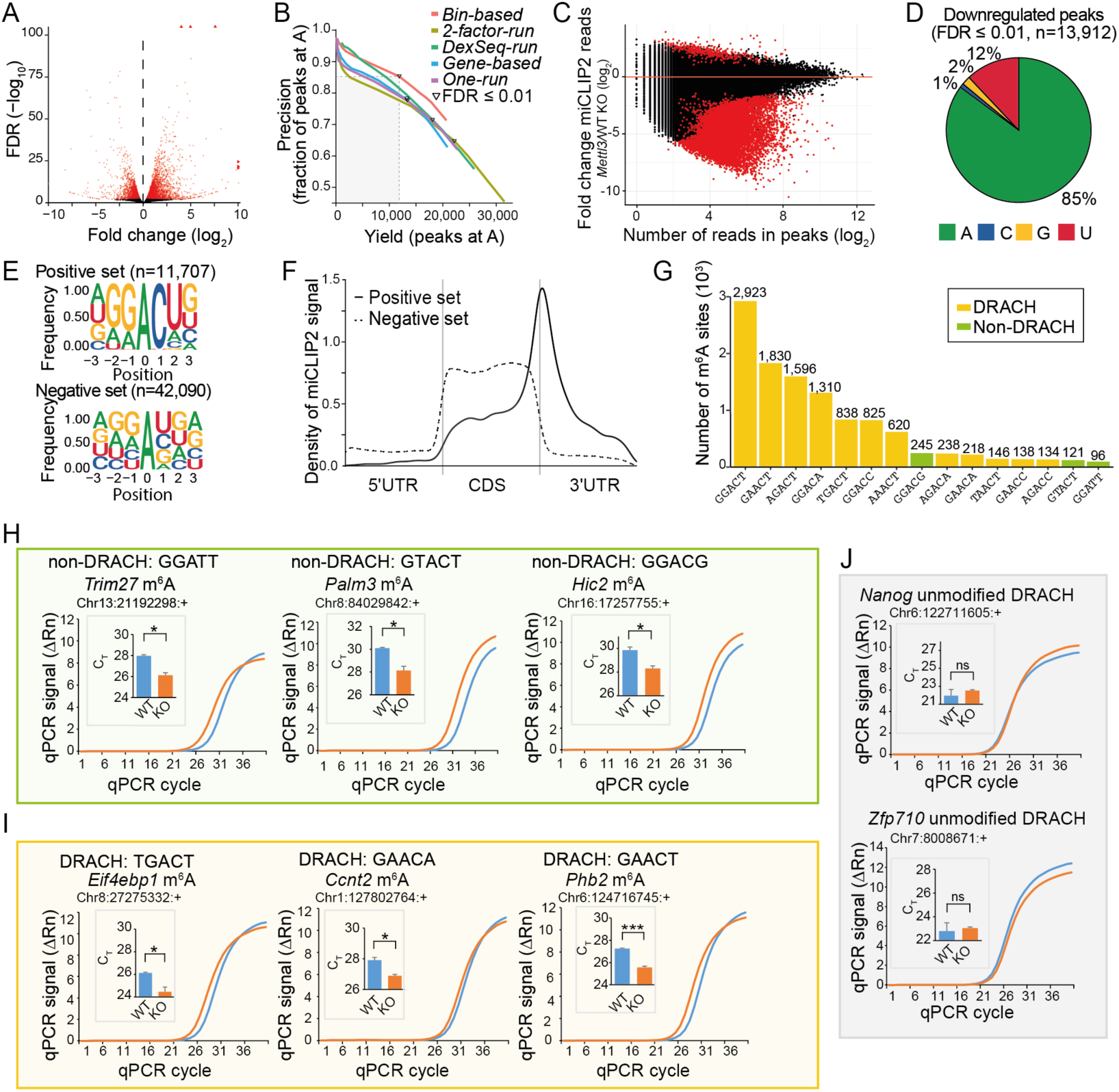
Differential peak analysis allows to identify true m^6^A sites from miCLIP2 data. ***A.*** *Mettl3* KO causes drastic changes in gene expression. Volcano plot shows log_2_-transformed fold change (log2FC) of gene expression between WT and *Mettl3* KO against log_10_-transformed false discovery rate (FDR). Significantly changing genes are highlighted in red (FDR ≤ 0.01). ***B.*** The *bin-based* approach for differential methylation analysis outperforms other tested strategies. Number of identified peaks at A (x-axis) and fraction of peaks at A (y-axis) are given for different approaches (see **Supplemental Material**, Section A). Curves were generated by step-wise increases in stringency (FDR). FDR ≤ 0.01 is marked for each approach. ***C.*** Most changing peaks go down upon *Mettl3* KO. Comparison of log2FC in miCLIP2 signal per peak between WT and *Mettl3* KO (y-axis) against reads per peak (log_2_-transformed, x-axis). Significantly regulated peaks are highlighted in red (|log2FC| > 1, FDR ≤ 0.01). ***D.*** Most significantly downregulated peaks are located at adenosines. Pie chart represents nucleotide distribution of downregulated peaks. ***E.*** Sequence motifs of peaks in the positive (top) and negative (bottom) set. Logos show relative frequency of nucleotides at positions −3 to +3 around central A. ***F.*** Peaks in the positive set accumulate around stop codons. Density plot shows distribution of peaks in scaled transcript regions. UTR, untranslated region, CDS, coding sequence. ***G.*** The most frequent pentamers include non-DRACH motifs. Number of peaks (positive set) located at specific pentamer at DRACH (orange) and non-DRACH (olive) motifs. ***H-J.*** Selected m^6^A sites were validated by SELECT experiments. Exemplary real-time fluorescence amplification curves (normalised reporter value, ΔRn) and quantifications of threshold cycle (C_T_) values (technical replicates) for mESC WT versus *Mettl3* KO samples are shown for m^6^A sites at non-DRACH (H) and DRACH (I) motifs as well as unmodified DRACH motifs with a miCLIP2 peak (J). Neighbouring unmodified A nucleotides as control for each tested site are given in **Supplementary Figure S3D**. *** *P* value < 0.001, * *P* < 0.05, ns, not significant, two-sided Student’s *t*-test, n=3.

Among the DRACH motifs identified in the positive set, the most frequent pentamer was GGACT, followed by GAACT and AGACT (17) (**Figure 2G**). Surprisingly, however, we also detected 741 m^6^A sites (6.3%) at non-DRACH motifs (non-DRACH m^6^A). While most of these non-DRACH motifs still contained the AC dinucleotide (52), some also diverged from this, such as GGATT (**Figure 2G**). We used SELECT (single-base elongation- and ligation-based qPCR amplification) (29) as an orthogonal antibody-independent m^6^A detection method to test the reliability of our approach. To this end, we compared SELECT qPCR amplification curves from WT versus *Mettl3* KO samples for an exemplary non-DRACH m^6^A site from the positive set, located in the last exon of the *Trim27* gene (A at position chr13:21192298:+, GGATT). Indeed, we detected Mettl3-dependent methylation at A in the GGATT motif, reflected in a reduced efficiency of the qPCR amplification when the m^6^A mark is present (**Figure 2H**). As a control, we tested an adjacent A in the same gene (position chr13:21192294:+), which remained unchanged upon *Mettl3* KO (**Supplementary Figure S3D**). We similarly validated two out of two additional non-DRACH m^6^A sites in the genes *Palm3* (chr8:84029842:+, GTACT) and *Hic2* (chr16:17257755:+, GGACG) (**Figure 2H and Supplementary Figure S3D**). For comparison, we also confirmed three out of three m^6^A sites at *bona fide* DRACH motifs in the genes *Eif4ebp1* (chr8:27275332:+, TGACT), *Ccnt2* (chr1:127802764:+, GAACA) and *Phb2* (chr6:124716745:+, GAACT) (**Figure 2I and Supplementary Figure S3D).**

DRACH motifs were also present at 1,043 peaks (2.5%) in the negative set. The miCLIP2 signal at these peaks did not decrease in the *Mettl3* KO, indicating that the antibody may show a residual background activity against the DRACH motif itself. SELECT experiments for two out of two selected sites in the genes *Nanog* (chr6:122711605:+) and *Zfp710* (chr7:8008671:+) confirmed that the respective A indeed did not carry an N6-methyl modification (**Figure 2J**).

All together, we defined a positive set of more than 10,000 m^6^A sites, that are modified in a Mettl3-dependent manner. In addition to canonical DRACH motifs, we identified a fraction of m^6^A modifications at non-DRACH motifs which show the same characteristics and Mettl3 dependency as m^6^A sites at DRACH motifs.

### Machine learning allows to reliably predict m^6^A sites from miCLIP2 data

To allow for m^6^A detection independently of an accompanying KO dataset, we built a predictive machine learning model to discriminate true m^6^A sites from background signal in the miCLIP2 data (**Figure 3A**). For model training, we combined the positive (n=11,707) and negative (n=42,090) set identified in the differential methylation analysis upon *Mettl3* KO. The unbalanced setup was chosen to reflect the predominance of nonspecific background in the miCLIP2 data (**Figure 1D-G**). We randomly split the data into a training set (80%) and an independent test set (20%). The input variables for training included 10-nt flanking nucleotide sequence to either side of A, the transcript region and the relative signal strength. We further added, as orthogonal information, the number of coinciding C-to-T transitions in the read-through reads, which we initially removed from the data (**Figure 3B**, see **Supplementary Material**, Section B).

**Figure 3.**
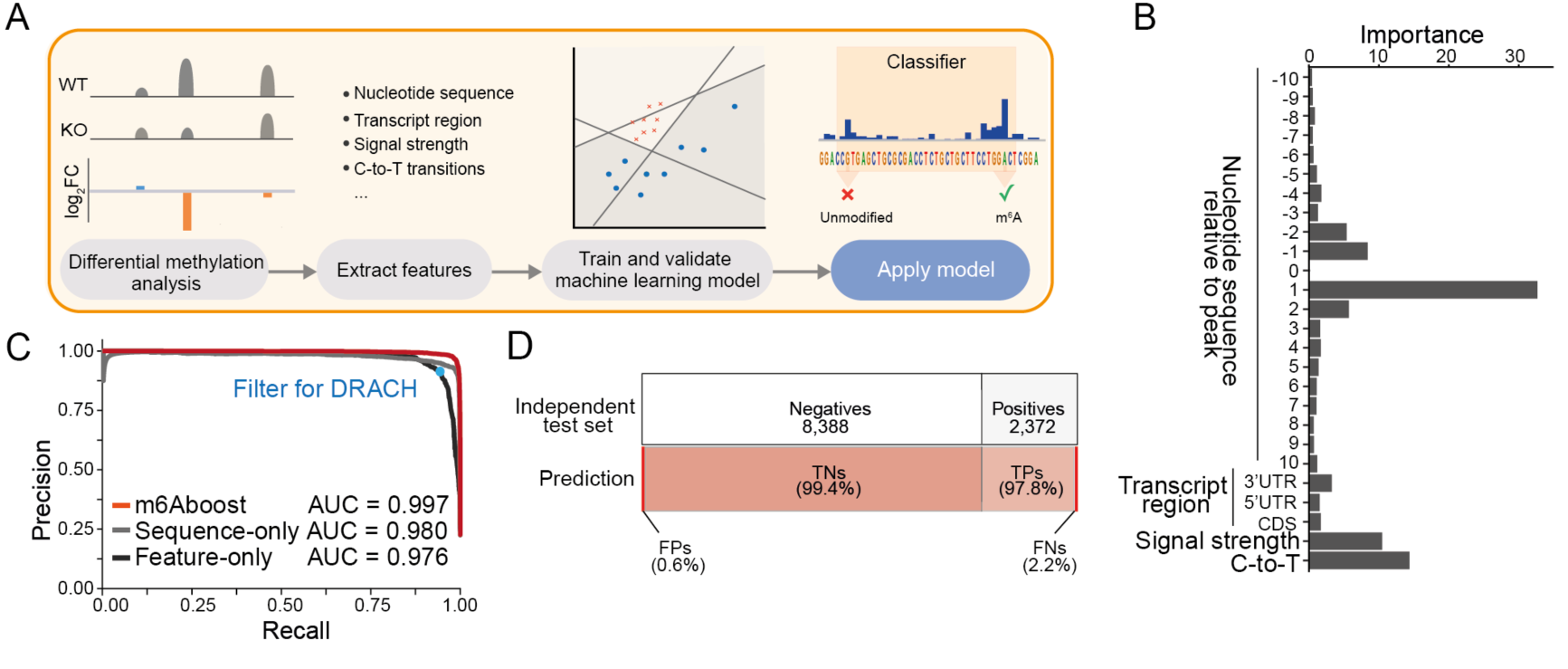
The machine learning classifier m6Aboost reliably predicts m^6^A sites from miCLIP2 data. ***A.*** Overview of building the machine learning classifier. First, miCLIP2 WT and *Mettl3* KO datasets are analysed for differential methylation to identify Mettl3-dependent m^6^A sites. The resulting positive and negative sets are used to extract features and train a machine learning model. The model is validated on an independent test set. Finally, the model can be applied to new miCLIP2 datasets to classify the miCLIP2 peaks as modified m^6^A sites versus unmodified background signal. ***B.*** Highest informative content lies in the nucleotide sequence, the relative signal strength of the peak and the number of C-to-T transitions. Bar plot shows the features used for m6Aboost prediction and their associated importance ranking. UTR, untranslated region, CDS, coding sequence. ***C.*** m6Aboost outperforms baseline models trained only on sequence (sequence-only) or experimental features (feature-only). Precision-recall curve shows performance of m6Aboost compared to baseline models with the corresponding area under the curve (AUC). Precision and recall when solely filtering for DRACH motifs is shown for comparison (blue dot). ***D.*** m6Aboost achieves 99% accuracy on an independent test set. Bars visualise composition of independent test set (n=10,760) from positive (22%) and negative (78%) peaks (top) and the resulting m6Aboost predictions (bottom). In total, 10,658 peaks (99%) were correctly predicted, while 102 peaks were misclassified. TNs, true negatives, TPs, true positives, FNs, false negatives, FPs, false positives.

We tested three different machine learning algorithms, which consistently reached high predictive accuracy (support vector machine, random forest, and adaptive boosting [AdaBoost]; **Supplementary Figure S4A-E**, see **Supplementary Material**, Section B). Following a series of benchmarks, we chose the AdaBoost-based predictor, which we named m6Aboost. AdaBoost is a boosting ensemble algorithm that weights the input for each iteration by the misclassification errors from previous iterations, and thereby improves the accuracy of the final predictions (53). The error rate of m6Aboost on the independent test set reached 0.99%, with > 99% area under the curve (AUC) in a precision-recall curve (**Figure 3C and Supplementary Figure S4A and D**). Evaluation on an independent test set showed that 99% of sites were correctly classified (**Figure 3D**). The performance was confirmed by five-fold cross-validation (**Supplementary Figure S4C**). The highest informative content was attributed to the immediate sequence around the modified A nucleotide, the relative signal intensity of peaks, and orthogonal information on C-to-T transitions (**Figure 3B**). Baseline models trained only on sequence information (position −10 to +10; ‘sequence-only’) or experimental features (relative signal strength, C-to-T transitions, and transcript region; ‘feature-only’) achieved worse classification results (**Figure 3C**), supporting that both types of features are required for optimal performance. Consistently, our m6Aboost outperformed a simple filter for DRACH motifs (**Figure 3C**, blue dot).

### m6Aboost predicts m^6^A sites also in lowly expressed transcripts

To test the algorithm on a complete miCLIP2 dataset, we applied m6Aboost to all peaks on A nucleotides in the mESC WT miCLIP2 data (n=117,142). In total, m6Aboost extracted 25,456 putative m^6^A sites in 9,363 genes (**Figure 4A**). These included 11,548 sites from our initial positive set (98.6% of positive set) plus 13,908 additional m^6^A sites. The latter were enriched in lowly expressed genes and most likely failed to reach significance in the differential methylation analysis due to low read counts (**Supplementary Figure S4F**). The miCLIP2 signal in all sites coherently went down in the *Mettl3* KO (94% with log2FC < −1; **Figure 4B**), supporting that they are indeed true m^6^A sites.

**Figure 4.**
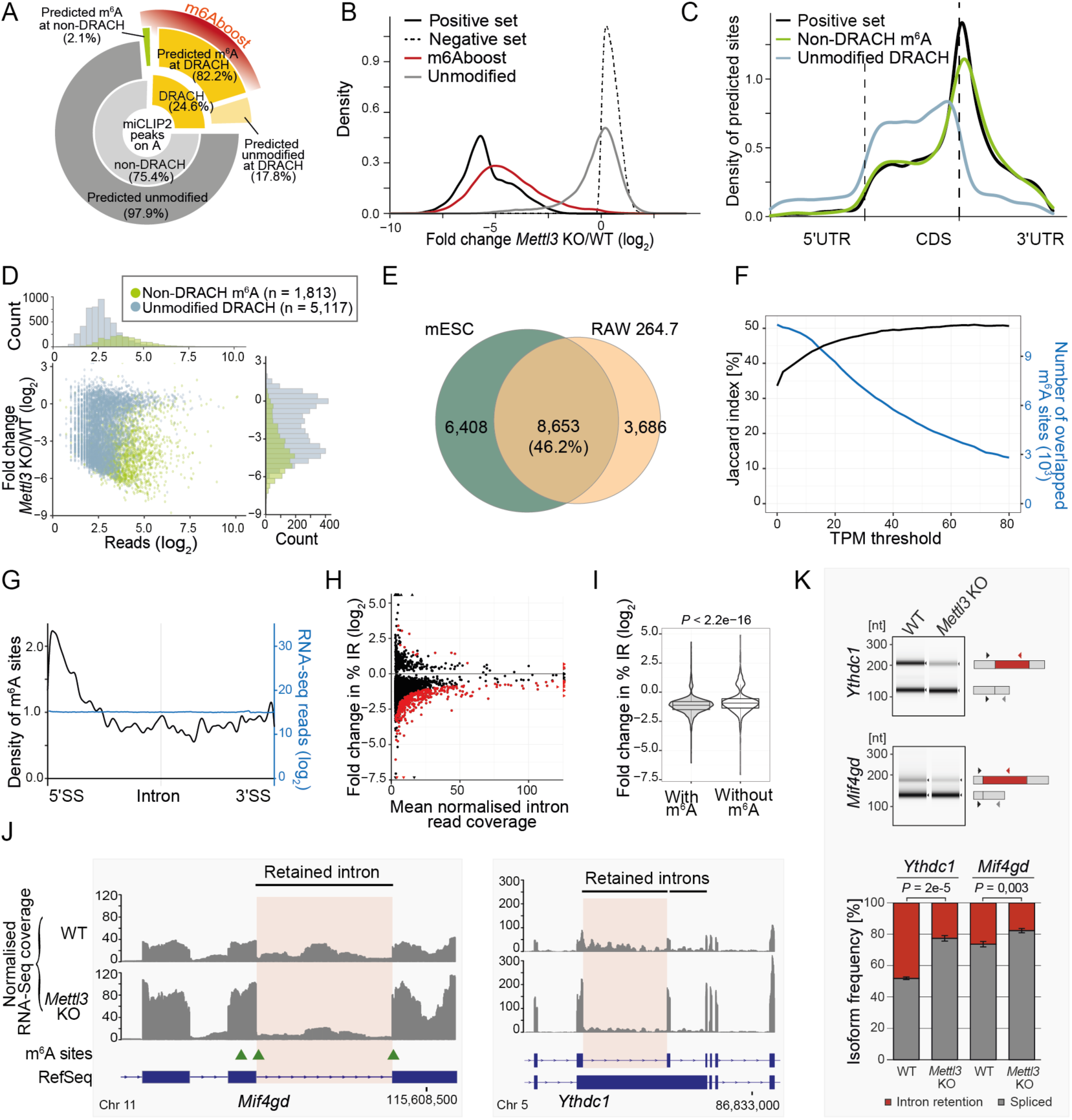
m^6^A sites occur at non-DRACH motifs and accumulate in retained introns. ***A.*** m6Aboost predicts m^6^A sites at DRACH and non-DRACH motifs in mESC WT miCLIP2 data. Inner circle of donut chart shows occurrence of DRACH (n=28,760, 24.6%) and non-DRACH (n=88,382, 75.4%) motifs for all miCLIP2 peaks at A. Outer circle shows m6Aboost prediction results (marked in red) with 23,643 m^6^A sites and 5,117 unmodified sites at DRACH (82.2% and 17.8%, respectively, of all peaks at DRACH) as well as 1,813 m^6^A sites and 86,569 unmodified sites at non-DRACH (2.1% and 97.9%, respectively, of all peaks at non-DRACH). ***B.*** Predicted m^6^A sites (n=25,456) go down upon *Mettl3* KO, whereas predicted unmodified sites (n=91,686) remain unchanged. Density plot shows distribution of log2-transformed fold changes in miCLIP2 signal between *Mettl3* KO and WT samples. Positive and negative set are shown for comparison. ***C.*** m^6^A sites at non-DRACH motifs (n=1,813) show a similar accumulation at stop codons as the positive set. Visualisation as in Figure 2F. ***D.*** m6Aboost predicts that not all peaks at DRACH motifs are m^6^A sites. Scatter plot and histograms show fold change in miCLIP2 signal (log2-transformed, y-axis) against number of reads per peak (log_2_-transformed, x-axis) for 5,117 peaks at DRACH motifs (light blue) that are predicted to be unmodified by m6Aboost. m^6^A sites at non-DRACH motifs (olive) are shown for comparison. ***E.*** Most m^6^A sites are shared between two mouse cell lines. Venn diagram shows overlap of predicted m^6^A sites in expressed genes (TPM ≥ 20, n=4,490) from mESC WT and RAW 264.7 cells. Venn diagram without expression filter is shown in **Supplementary Figure S5E**. ***F.*** Overlap of m^6^A sites between two mouse cell lines increases in higher expressed genes. TPM threshold representing the gene expression (x-axis) against the Jaccard index (y-axis). Number of overlapping m^6^A sites are shown as comparison (blue). ***G.*** m^6^A sites accumulate towards the 5’ splice sites of introns. Metaprofile shows density of m^6^A sites along scaled introns (n=3,509 m^6^A sites on 1,465 different Introns). Coverage of RNA-seq reads on the same introns is shown for comparison (blue). SS, splice site. ***H.*** Intron retention (IR) is globally reduced in the *Mettl3* KO cells. Scatter plot shows fold change in relative IR (%IR, y-axis) against mean normalised RNA-seq reads on the introns across all samples (x-axis) for 4,925 measured IR events. 401 significantly changed IR events are highlighted in red (FDR ≤ 0.05). ***I.*** Introns harbouring m^6^A sites show a significant trend towards IR reduction. Violin plot compares fold changes in %IR for retained introns with (n=4,098) and without (n=827) m^6^A sites. *P* value < 2.22e-16, Wilcoxon rank-sum test. ***J.*** IR is reduced in the *Mif4gd* and *Ythdc1* transcripts. Genome browser view of RPGC-normalised RNA-seq coverage are shown for merged replicates from WT and *Mettl3* KO mESCs. Predicted m^6^A sites are indicated with green arrowheads. IR events validated in (K) are highlighted in red. ***K*.** Frequency of *Ythdc1* and *Mif4gd* IR isoforms is lower in *Mettl3* KO mESCs. Semiquantitative three-primer RT-PCR to quantify isoform frequencies in WT and *Mettl3* KO cells, with shared forward and isoform-specific reverse primers displayed next to corresponding PCR products in capillary gel electrophoresis (top). Quantification of relative band intensities (bottom) is displayed as mean ± s.d.m., n=3, unpaired two-sided Student’s *t*-test.

Of note, 1,813 out of 25,456 (7.1%) predicted m^6^A sites resided at non-DRACH motifs (**Figure 4A**). These non-DRACH m^6^A sites showed an enrichment nearby stop codons similar to the positive set and the vast majority were depleted in the *Mettl3* KO (**Figure 4C and D**), supporting that predicted non-DRACH sites are indeed true m^6^A sites. On the other hand, m6Aboost predicted that not all peaks at DRACH motifs corresponded to true m^6^A sites. Indeed, about half of these sites did not respond to *Mettl3* KO and distributed similarly to the negative set (**Figure 4D and Supplementary Figure S4G**), suggesting that the m^6^A-specific antibody shows a residual activity towards the unmodified DRACH motif. The other half had low read counts and preferentially resided in lowly expressed genes (**Supplementary Figure S4G**), possibly leading to their misclassification. Importantly, m6Aboost associates a prediction score with each site that allows to minimise the number of false positives, at the expense of false negatives, by tightening the prediction score threshold (**Supplementary Figure S4H and I**). Altogether, we conclude that m6Aboost efficiently discriminates relevant signal from nonspecific background, offering a reliable prediction of genuine m^6^A sites from miCLIP2 data.

As an orthogonal support, we compared our predicted m^6^A sites to those detected by the antibody-independent method MAZTER-seq in the same cell line (21). MAZTER-seq relies on the methylation-sensitive RNase MazF which cleaves at unmethylated ACA motifs. We found that 34.5% of the reliably identified m^6^A sites from MAZTER-seq (200 out of 580 sites) were also present in our data, further supporting the validity of our approach.

For comparison, we also performed miCLIP2 experiments on poly(A)+ RNA from RAW 264.7 cells, a mouse macrophage cell line (three biological replicates, 29.8 million truncation events on average). Out of 462,073 miCLIP2 peaks, m6Aboost identified a total of 19,301 m^6^A sites (**Supplementary Table S1**). Overlay with the mESC data showed that a third of the predicted m^6^A sites were shared between both cell lines, rising to about 50% when focussing on genes that were highly expressed in both cell lines (TPM ≥ 20 or more; **Figure 4E and F and Supplementary Figure S5E**).

### m^6^A depletion triggers efficient splicing of retained introns

Since our miCLIP2 data was generated for poly(A)-selected RNA, most identified m^6^A sites were located in exons. However, we also detected a number of m^6^A sites in retained introns. Interestingly, the intronic m^6^A sites showed a strong accumulation towards the 5’ splice sites (**Figure 4G**), suggesting that they might impact intron splicing. Indeed, using IRFinder (47), we could identify 401 significantly changed intron retention (IR) events in the RNA-seq data of *Mettl3* KO mESCs (change in IR [|ΔIR|] > 3%, FDR < 0.05; **Figure 4H and I**). 384 out of 401 significantly changed introns showed reduced coverage in the *Mettl3* KO, as seen for intron 5 in *Mif4gd* and intron 11 in *Ythdc1* (**Figure 4J**), indicating increased splicing efficiency. Isoform-specific semiquantitative RT-PCR confirmed a lower frequency of the *Ythdc1* and *Mif4gd* isoforms with retained introns in *Mettl3* KO mESCs (**Figure 4K**). This trend was also reflected in a global reduction in IR across the transcriptome, as 4,563 out of 4,925 measured IR events (92.7%) showed a ΔIR < 0 (**Figure 4H**). Generally, introns harbouring m^6^A modifications showed a significant trend towards more IR reduction compared to unmodified introns (**Figure 4I**), indicating that modifications on retained introns may directly influence splicing efficiency.

### m6Aboost can be applied to predict m^6^A sites in human cells

To test m6Aboost on miCLIP2 data from a different species, we performed miCLIP2 experiments with poly(A)+ RNA from human HEK293T cells (n=4 replicates with 30 million truncation events on average, **Supplementary Figure S1F and G**). Starting from more than 788,758 miCLIP2 peaks, m6Aboost identified 36,556 m^6^A sites in 7,552 genes, corresponding to 21% of all peaks at A (**Supplementary Table S1**). The m^6^A sites occurred with a median of three sites per gene and accumulated around stop codons (**Figure 5A and Supplementary Figure S5A**), mirroring the distribution in the mouse cells.

**Figure 5.**
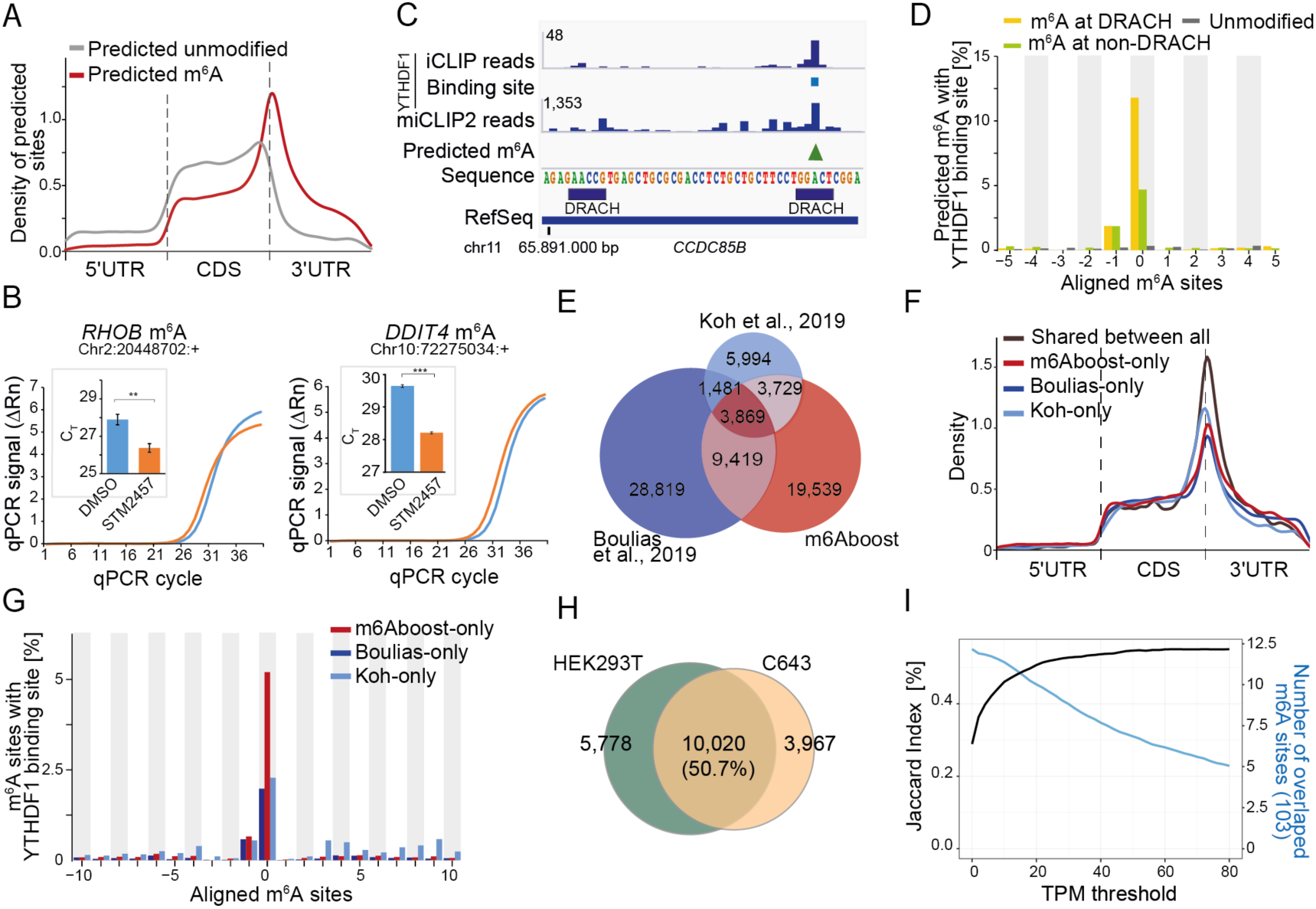
m6Aboost predicts 36,556 m^6^A sites from HEK293T miCLIP2 data. ***A.*** Predicted m^6^A sites are located around the stop codon. Visualisation as in Figure 2F. ***B.*** Selected m^6^A sites were validated by SELECT with HEK293T cells treated with METTL3 inhibitor (STM2457) or DMSO control. Visualisation as in Figure 2H. Neighbouring unmodified A nucleotides as control are given in **Supplementary Figure S5C**. *** *P* value < 0.001, ** *P* < 0.01, two-sided Student’s *t*-test, n=3. ***C.*** Predicted m^6^A sites overlap with YTHDF1 binding sites. Genome browser view of the gene *CCDC85B* shows crosslink events from published YTHDF1 iCLIP data for HEK293T together with miCLIP2 signal (merge of four replicates) and m6Aboost-predicted m^6^A site (green arrowhead) from our HEK293T miCLIP2 data. ***D.*** YTHDF1 precisely binds at the predicted m^6^A sites. Percentage of m^6^A sites (position 0) with YTHDF1 binding sites (y-axis) in a 21-nt window are given for predicted m^6^A sites at DRACH (yellow) and non-DRACH (green), as well as predicted unmodified sites (grey). ***E.*** Predicted m^6^A sites from HEK293T miCLIP2 overlap with published m^6^A data. Venn diagram shows single-nucleotide overlap with miCLIP and m6ACE-seq data (m^6^A antibody by Synaptic Systems and Abcam, respectively). Note that m^6^A sites in Boulias et al, 2019 had been filtered for DRACH motifs. ***F,G.*** Analysis of m^6^A sites that are unique to one of the three datasets compared in (E). ***F.*** Unique m^6^A sites accumulate around stop codons. Visualisation as in Figure 2F. ***G.*** Unique m^6^A sites are enriched in YTHDF1 binding sites. Visualisation as in (D). ***H.*** Most m^6^A sites are shared between two different human cell lines. Venn diagram shows overlap of predicted m^6^A sites in expressed genes (TPM ≥ 20, n=3,298) from HEK293T and C643 cells. A Venn diagram without expression filter is shown in **Supplementary Figure S5E**. ***I.*** More m^6^A sites are shared between two different mouse cell lines in higher expressed genes. TPM threshold representing the gene expression (x-axis) against the Jaccard index (y-axis). Number of overlapping m^6^A sites are shown as comparison (blue).

We used SELECT to validate the presence of m^6^A modifications in HEK293T cells in an antibody-independent manner (29). In order to deplete m^6^A, we employed a specific METTL3 inhibitor (STM2457, STORM Therapeutics) (28), which progressively reduced the relative m^6^A levels with increasing concentration, down to 22% (**Supplementary Figure S5B**). We then compared SELECT qPCR amplification curves from inhibitor-treated HEK293T cells against DMSO control samples for three exemplary m^6^A sites. This confirmed the presence of m^6^A in two out of three sites in the genes *DDIT4* (chr10:72275034:+) and *RHOB* (chr2:20448702:+) (**Figure 5B**). As a control, we tested adjacent A sites in the same genes which remained unchanged upon METTL3 inhibition (*DDIT4*: chr10:72275038:+; *RHOB*: chr2:20448698:+; **Supplementary Figure S5C**). A third putative m^6^A site could not be validated (*ABT1*: chr6:26598621:+).

As an independent line of evidence, we overlapped the m6Aboost-predicted m^6^A sites with binding sites of the cytoplasmic m^6^A reader protein YTHDF1 from published iCLIP data (54). Metaprofiles showed a sharp peak in YTHDF1 binding precisely at the predicted m^6^A sites at DRACH motifs (**Figure 5C and D and Supplementary Figure S5D**). Although less pronounced, we detected considerable YTHDF1 binding also at predicted m^6^A sites at non-DRACH motifs, further supporting that these indeed represent genuine m^6^A sites.

We compared our predicted m^6^A sites in HEK293T with published validated m^6^A sites in the same cell line by the antibody-independent method SCARLET that uses thin-layer chromatography (52). We found that all m^6^A sites with > 5% methylation in HEK293T cells were also present in our data, whereas sites that were not validated by SCARLET (< 5% methylation) were not detected by miCLIP2 (**Supplementary Table S3**). To further support the predicted m^6^A sites, we compared our miCLIP2 data with published miCLIP and m6ACE-seq data for the same cell line (51, 55). m6A-Crosslinking-Exonuclease-sequencing (m6ACE-seq)-seq is a recently developed tool which incorporates 5ʹ to 3ʹ exoribonuclease treatment after m^6^A-antibody crosslinking to increase the resolution and omit radioactive gel electrophoresis (55). We found that almost half of our m^6^A sites overlapped at single-nucleotide level with at least one further dataset (**Figure 5E**). The remaining sites occurred on lowly expressed genes, but still showed an m^6^A-typical distribution along transcripts and overlapped with YTHDF1 binding (**Figure 5F and G and Supplementary Figure S5F**). This suggests that these m^6^A sites were missed in other studies due to experimental variability and technical limitations rather than lack of modification.

As a second human cell line, we performed miCLIP2 experiments on poly(A)+ RNA from C643 cells, a human thyroid cancer cell line (three biological replicates, **Supplementary Table S1**). Here, m6Aboost predicted a total of 18,789 m^6^A sites. Comparison with HEK293T showed that similar to mouse, 50.7% of all m^6^A sites on highly expressed genes were shared between mESC and C643 cells (TPM ≥ 20 or higher; **Figure 5H and I and Supplementary Figure S5E**), an estimate that is stable with increasing expression. We therefore conclude that about half of all m^6^A modifications are constitutively present in different cell types in human and mouse.

### miCLIP2 allows to map m^6^A sites from low input material

Most current protocols for antibody-based m^6^A detection start from 5-10 μg of poly(A)+ mRNA (37, 56). In our standard setup, we use just 1 μg, from which we obtain more than 30 million unique miCLIP2 reads on average with low PCR duplication rates (**Supplementary Table S1**). However, when working with scarce material such as tissue samples, the amount of extractable RNA is often limited. We therefore tested whether miCLIP2 can be applied with even lower RNA input. To this end, we used poly(A)+ mRNA from mouse heart tissue samples and titrated the amount of input RNA down to 50 ng. The resulting miCLIP2 libraries contained 2-50 million truncation events (**Supplementary Table S1**).

We found that even with these small amounts of input RNA, the miCLIP2 signals were still reproducible at nucleotide level (**Figure 6A and Supplementary Figure S5G**). As expected, the sensitivity of miCLIP2 progressively decreased with lower input material. The precision, however, was hardly compromised, since the identified sites were highly overlapping at all concentrations (**Figure 6B**). Moreover, m^6^A sites from all RNA input concentrations were consistently enriched at DRACH motifs and nearby stop codons (**Figure 6C and D**). Together, these results suggest that our approach can be used to identify m^6^A modifications even from a limited amount of input RNA.

**Figure 6.**
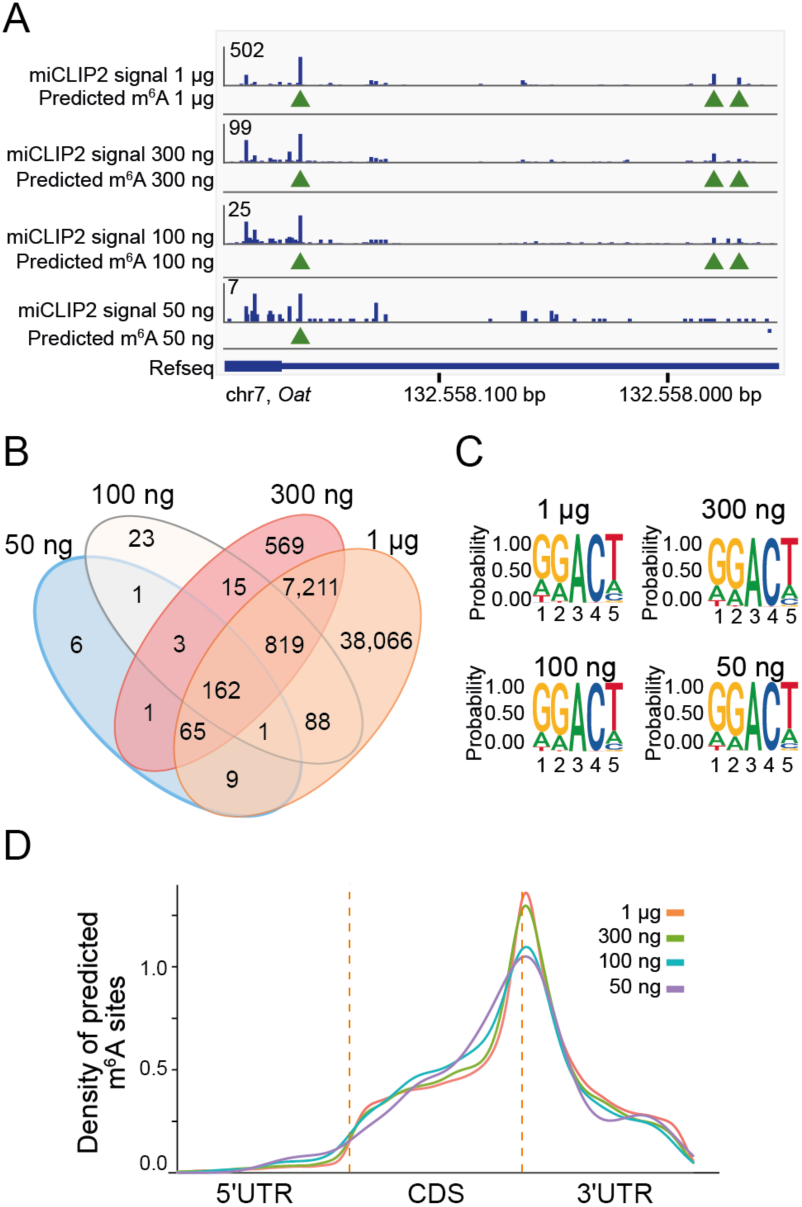
miCLIP2 allows to map m^6^A sites from low input material. ***A.*** m6Aboost predicts overlapping m^6^A sites from miCLIP2 data for different RNA input concentrations. Example genome browser view of the *Oat* gene shows miCLIP2 signals and corresponding m6Aboost predictions (green arrowheads) for 1 μg, 300 ng, 100 ng and 50 ng of input RNA. ***B.*** The majority of m^6^A sites predicted from low-input libraries overlap with 1 μg input library. Overview of the overlap of predicted m^6^A sites from different concentrations. ***C.*** All predicted sites from different concentrations resemble a DRACH motif. Sequence logo of the predicted m^6^A sites from miCLIP2 from different RNA input concentrations including the surrounding four nucleotides. ***D.*** Predicted m^6^A sites from miCLIP2 with different RNA input concentrations cluster around the stop codon. Visualisation as in Figure 2F.

## DISCUSSION

Knowledge on the precise location of m^6^A sites is essential to unravel the molecular effects and biological functions of this universal RNA modification. With the advent of next-generation sequencing, new experimental protocols allow for a systematic mapping of m^6^A sites, often with single-nucleotide resolution (57). Although alternative methods recently became available (21,22,58,59), the most widely used approaches rely on a set of available antibodies against the modified nucleotide (57). These methods suffer from the broad reactivity of these antibodies, which cross-react with unmodified adenosines or related modifications such as m^6^Am, thereby generating excess false positives (17). Moreover, many protocols require high amounts of starting material, or target only a restricted subset of m^6^A sites that occur for instance in a specific sequence context (21,22,37,56). In this study, we tackle these limitations by combining the optimised miCLIP2 protocol and the machine learning model m6Aboost to reliably map m^6^A modifications at high resolution and depth. Our approach builds on three major experimental and computational innovations that are critical for its efficiency and accuracy.

First, we improved the efficiency of the experimental miCLIP2 protocol by incorporating the recently published iCLIP2 library preparation (24), including separately ligated adapters, two rounds of PCR amplification and a bead-based clean-up strategy. This reduces the processing time to just four days and provides high-complexity datasets without PCR duplicates. With this setup, we now routinely obtain more than 30 million unique miCLIP2 reads from 1 μg input RNA - twenty times less than in the original protocol (17). The moderate duplication rate (**Supplementary Table S1**) indicates the miCLIP2 libraries in this study were not sequenced to saturation, suggesting that many more m^6^A sites could still be identified from the same libraries. Moreover, it is possible to obtain reproducible data down to 100 ng and less of input RNA. The reduced input requirement will be particularly useful for studies on nascent RNA or clinical samples and *in vivo* disease models where starting material is limiting.

Second, we tackled the high false positive rate from the m^6^A-specific antibodies, which is inherent to antibody-based approaches, through the direct comparison with *Mettl3* KO cells. Using a custom-tailored differential methylation analysis strategy, we identify more than 10,000 Mettl3-dependent m^6^A sites in the WT mESC miCLIP2 data that constitute the positive set of high-confidence m^6^A sites for subsequent model training (see below). Of note, we find that m^6^A modifications occur outside of DRACH motifs (6.3% of all predicted m^6^A sites) and validate selected m^6^A sites at non-DRACH motifs using an orthogonal antibody-independent method. Similar motifs were previously reported and recently confirmed in direct RNA sequencing data (Oxford Nanopore Technologies) (17, 60). Importantly, since the m^6^A sites at non-DRACH motifs were included in the m6Aboost model training, similar sites can be readily identified in future miCLIP2 experiments. In addition, we propose that the sequence composition of the high-confidence m^6^A sites from the differential methylation analysis (**Figure 2E**), captured for instance in a position weight matrix, could be used to filter other datasets in a more effective way. Moreover, our strategies to account for changes in transcript abundance in order to identify differentially methylated sites will be applicable for other RNA modifications, such as 5-methylcytosine (m^5^C) in m^5^C-miCLIP (57, 61).

Third, we trained a machine learning model, termed m6Aboost, to accurately extract Mettl3-dependent m^6^A sites from any miCLIP2 dataset. Several machine learning approaches have been developed to predict m^6^A sites from the primary RNA sequences (62–64). However, most existing models were trained on data of limited resolution and size, and consequently perform poorly for single-nucleotide predictions. Here, we apply machine learning to predict m^6^A sites in miCLIP2 data based on a high-confidence positive set of Mettl3-dependent m^6^A sites. We therefore tackle the inherent problem of false positives that impair most antibody-based m^6^A detection protocols (57). The resulting m6Aboost model allows one to transfer our gained knowledge to other miCLIP datasets without the need for an accompanying *Mettl3* KO, which is not feasible in many biological settings (27). Because m6Aboost allows for m^6^A sites at non-DRACH motifs and sorts out false positive miCLIP2 signals, even at DRACH, it outperforms the commonly used DRACH motif filter (37,51,59). The stringency against false positives can be tuned according to the requirements of the user by adjusting the prediction score of m6Aboost.

We note that our model was trained on miCLIP2 data that was obtained with a specific m^6^A antibody (Synaptic Systems). It is known that certain biochemical features such as the truncation rate at the crosslinked antibody and the distribution of C-to-T transitions varies with each antibody (17, 57). We envision that our approach can be retrained on data for other antibodies against m^6^A and other RNA modifications that can be mapped via miCLIP2, if an accompanying depletion dataset is available. This includes the related RNA modification m^6^Am, which is present in the miCLIP2 data due to cross-reactivity of the m^6^A antibody, and could be recognised and specifically discriminated from m^6^A after retraining upon depletion of the m^6^Am-specific methyltransferase PCIF1 (51, 65).

In this study, we generated m^6^A profiles for four human and mouse cell lines that will serve as a resource for future studies. Comparing the methylation profiles revealed that about half of all m^6^A sites are shared between cell lines in either species. Moreover, we confirm that m^6^A is mainly deposited around stop codons and within the 3’ UTR (15, 16). Interestingly, we also observe an accumulation near the 5’ splice sites of retained introns. Further, our data indicates that m^6^A can promote intron retention. Previous studies rather described an increase in intron retention events in *Mettl3* KO mESC cells (27), or in null mutants of the Mettl3 orthologue Ime4 in *Drosophila melanogaster* (4,66,67). In contrast, a recent study found that TARBP2-dependent m^6^A deposition in introns prevents splice factor recruitment and efficient intron excision (68), in line with our observations. This adds a new angle to the controversy surrounding the impact of m^6^A modifications on alternative splicing. While some studies reported on extensive splicing alterations upon *Mettl3* depletion, others rebutted a strong connection between m^6^A and splicing (69–72). Consistent with the latter view, we generally observe very few changes in cassette exon splicing in the *Mettl3* KO mESCs. Intron retention, however seemed to be systemically affected, with retained introns being spliced more efficiently throughout the transcriptome of *Mettl3* KO cells.

In essence, the combination of miCLIP2 and m6Aboost allows for a deep and accurate detection of m^6^A sites. Our study illustrates how artificial intelligence helps to eliminate background signals in order to decode high-throughput data and thereby aids to improve the precise analysis of m^6^A sites with nucleotide resolution.

## AVAILABILITY

The computational code is available in the GitHub repository (https://github.com/ZarnackGroup/Publications/tree/main/Koertel_et_al_2021).

## ACCESSION NUMBERS

All miCLIP2 and RNA-seq data sets generated in this study were submitted to the Gene Expression Omnibus (GEO) under the SuperSeries accession number GSE163500 (https://www.ncbi.nlm.nih.gov/geo/query/acc.cgi?&acc=GSE163500).

## SUPPLEMENTARY DATA

Supplementary Data are available at NAR online.

## ACKNOWLEDGEMENT

We gratefully acknowledge Tina Lence and Jean-Yves Roignant for help with miCLIP, Lina Worpenberg with initial help with SELECT, Yannic Schumacher for preparing poly(A)^+^ RNA from RAW 264.7 macrophages and Eric Miska for discussion. We are grateful to Marcel Schulz for advice on machine learning. Support by the IMB Genomics Core Facility and the use of its NextSeq500 (INST 247/870-1FUGG) are gratefully acknowledged. We kindly thank members of the IMB Genomics and Bioinformatics Core Facilities for technical assistance and reagents as well as all members of the König and Zarnack groups for lively discussions. NK, CR and MP were supported by the International PhD Programme on Gene Regulation, Epigenetics & Genome Stability, Mainz, Germany.

